# The entorhinal cortical alvear pathway differentially excites interneuron subtypes in hippocampal CA1

**DOI:** 10.1101/2020.01.22.915876

**Authors:** Karen A. Bell, Rayne Delong, Priyodarshan Goswamee, A. Rory McQuiston

## Abstract

The entorhinal cortex alvear pathway is a major excitatory input to hippocampal CA1, yet nothing is known about its physiological impact. We investigated the alvear pathway projection and innervation of neurons in CA1 using optogenetics and whole cell patch clamp methods in transgenic mouse brain slices. Using this approach, we show that the medial entorhinal cortical alvear inputs onto both CA1 pyramidal cells and stratum oriens interneurons were monosynaptic, had low release probability, and were mediated by AMPA receptors. Optogenetic theta burst stimulation was unable to elicit suprathreshold activation of CA1 pyramidal neurons but was capable of activating CA1 stratum oriens interneurons. CA1 stratum oriens interneuron subtypes were not equally affected. Higher burst action potential frequencies were observed in parvalbumin-expressing interneurons relative to vasoactive-intestinal peptide-expressing or a subset of oriens lacunosum-moleculare interneurons. Furthermore, alvear excitatory synaptic responses were observed in greater than 70% of PV and VIP interneurons and less than 20% of O-LM cells. Finally, greater than 50% of theta burst-driven inhibitory postsynaptic current amplitudes in CA1 PCs were inhibited by optogenetic suppression of PV interneurons. Therefore, our data suggest that the alvear pathway primarily affects hippocampal CA1 function through feedforward inhibition of select interneuron subtypes.

## Introduction

The hippocampus receives the majority of its excitatory input from the entorhinal cortex (EC). The EC projects to hippocampal CA1 via two converging pathways. One is the temporo-ammonic (TA) pathway that perforates the subiculum and courses through the stratum lacunosum-moleculare (SLM) to synapse on CA1 pyramidal cell (PC) and interneuron dendrites located in the SLM (Steward O 1976; Steward O and SA Scoville 1976). The other is the alvear pathway that courses through the alveus and adjacent stratum oriens (SO) of hippocampal CA1. After coursing transversely through the alveus, EC axons of the alvear pathway turn to ascend through all layers of hippocampal CA1 to terminate on PC and interneuron dendrites in the SLM (Deller T et al. 1996). Although the alvear pathway ultimately terminates in the SLM, alvear pathway axons in the SO have synaptic boutons that are thought to form excitatory synaptic connections with CA1 PC and interneuron dendrites (Deller T *et al*. 1996). However, the physiological impact of the synapses in the SO has not been investigated.

It is well known that the EC and hippocampus have crucial roles in the formation of long-term memories. In rodent, the alvear pathway carries most of the entorhinal input to dorsal hippocampal CA1 (Deller T *et al*. 1996). In humans, the alvear pathway also plays a prominent role in relaying inputs from the EC to hippocampal CA1 (Zeineh MM et al. 2017). In addition to being important for normal brain function, the EC is one of the first regions in the brain to display neurodegeneration and dysfunction in Alzheimer’s disease (AD) (Braak H and K Del Trecidi 2015). The initial pathology associated with AD is hallmarked by an accumulation of hyperphosphorylated tau protein that misfolds and eventually forms insoluble intracellular neurofibrillary tangles in EC projection neurons (Braak H and E Braak 1991, 1992; Braak H et al. 2006; Braak H and K Del Trecidi 2015). Tau spreads trans-synaptically from the EC to hippocampal CA1 (stage II) and from there to other parts of the cerebrum (stages III-VI). Significantly, in AD patients, tau protein has been shown to accumulate in the terminals of the alvear pathway but not the perforant path (Shukla C and LR Bridges 2001), suggesting that the alvear input may be particularly vulnerable at early stages of the disease. Despite the importance of the alvear pathway in normal and pathological states, the physiological effect of the alvear pathway on the postsynaptic neurons and network of hippocampal CA1 has not, to our knowledge, been reported.

However, the effect of EC stimulation on the excitability of hippocampal CA1 has been studied extensively. Some groups have observed a monosynaptic suprathreshold activation of CA1 PCs following electrical stimulation of the EC *in vivo* (Segal M 1972; Yeckel MF and TW Berger 1990, 1995), whereas other groups observed only subthreshold events (Andersen P et al. 1966; Leung LS et al. 1995). In contrast, *ex vivo* studies have demonstrated that the primary effect of stimulating hippocampal CA1 TA inputs (Empson RM and U Heinemann 1995, 1995) or all afferents in the SLM (including the TA, alvear, and thalamic nucleus reuniens pathways) is feedforward inhibition of hippocampal CA1 PCs (Colbert CM and WB Levy 1992; Dvorak-Carbone H and EM Schuman 1999; Ang CW et al. 2005). Notably, it appears that specific subsets of interneurons can be potently activated by SLM excitatory inputs, whereas others are only weakly effected or not effected at all (Milstein AD et al. 2015). However, the alvear inputs are in an anatomical distinct position to engage unique groups of interneurons in hippocampal CA1 and may have a differential effect on hippocampal network function relative to the TA input. To explore this, we expressed the optogenetic excitatory protein oChIEF in layer 3 (L3) of medial entorhinal cortex (MEC) to study the impact of these inputs onto PC and interneurons in the SO of hippocampal CA1. The MEC was chosen as the viral expression target by virtue of the transgenic mice available (see Materials and Methods). We found that activation of the alvear pathway produced subthreshold excitatory postsynaptic potentials (EPSPs) in CA1 PCs. In contrast, alvear pathway stimulation resulted in suprathreshold excitation of CA1 SO interneurons that in turn produced GABA_A_ inhibitory postsynaptic responses in CA1 PCs. A significant proportion of the interneurons activated by the alvear pathway were parvalbumin (PV)-expressing perisomatic interneurons that have been previously reported to have limited responses to TA pathway stimulation (Milstein AD *et al*. 2015). Therefore, our data suggest that, relative to the TA pathway, the alvear pathway can engage different hippocampal CA1 interneuron networks and may contribute to the generation of rhythms and perisomatic inhibition in hippocampal CA1.

## Materials and Methods

### Animals

NOP-tTA (B6.Cg-Tg(Klk8-tTA)QMmay/MullMmmh, MMRRC Stock No. 031780-MU) (Yasuda M and MR Mayford 2006), pOXR-1-Cre (C57BL/6N-Tg(Oxr1-cre)C14Stl/J, JAX Stock No. 030484) (Suh J et al. 2011), VIP-cre (*Vip^tm1(cre)Zjh^*/J, JAX Stock No. 010908) (Taniguchi H et al. 2011), PV-Cre (B6;129P2-*Pvalb^tm1(cre)Arbr^*/J, Jax Stock No. 008069) (Hippenmeyer S et al. 2005), PV-tdTomato (C57BL/6-Tg(Pvalb-tdTomato)15Gfng/J, JAX No. 027395) (Kaiser T et al. 2016), GIN (FVB-Tg(GadGFP)45704Swn/J) (Oliva AA, Jr. et al. 2000), Arch-GFP (B6;129*S-Gt(ROSA)26Sor^tm35.1(CAG-aop3/GFP)Hze^*/J, Jax Stock No. 012735) (Chow BY et al. 2010) and tetO-ChIEF-Citrine (Cheetham CEJ et al. 2016) mice used in these studies were housed in an animal care facility approved by the American Association for the Accreditation of Laboratory Animal Care. Experimental procedures followed the protocol approved by the Institutional Animal Care and Use Committee of Virginia Commonwealth University (AD20205). This protocol adhered to the ethical guidelines described in The Care and Use of Laboratory Animals 8^th^ Edition (Garber, 2011). All efforts were made to minimize animal suffering and to reduce the number of animals used.

### Breeding strategies

To examine layer 3 medial entorhinal cortex inputs to hippocampal CA1, we expressed the excitatory optogenetic protein oChIEF-mCitrine (Lin JY et al. 2009) in L3 MEC projection neurons using two different approaches. In the first approach, we injected the right hemisphere MEC of pOXR1-Cre mice with an AAV that expressed oChIEF-mCitrine in a Cre-dependent manner (rAAV-hSyn-Flex-oChIEF-mCitrine). In the second approach, we crossed the tTA-driver mouse line NOP-tTA (Yasuda M and MR Mayford 2006) with a tTA reporter mouse line tetO-ChIEF-Citrine (Cheetham CEJ *et al*. 2016) to produce NOP-tTA;tetO-ChIEF-Citrine mice. This cross resulted in the expression of oChIEF-mCitrine in layer 2 and L3 MEC projection neurons. We also expressed tdTomato or GFP selectively in interneuron subclasses to target interneuron subtypes for whole cell patch clamp recordings in hippocampal CA1 brain slices. To target PV interneurons, we crossed NOP-tTA;tetO-ChIEF-Citrine or pOXR1-Cre mice with either a homozygous cross of PV-Cre (Hippenmeyer S *et al*. 2005) and Ai14 (tdTomato reporter) (Madisen L et al. 2010) or a PV interneuron tdTomato expressing transgenic mouse line (PV-tdTomato) (Kaiser T *et al*. 2016). To target VIP interneurons, we crossed NOP-tTA;tetO-ChIEF-Citrine or pOXR1-Cre mice with a homozygous cross of VIP-Cre (Taniguchi H *et al*. 2011) and Ai14. Finally, we crossed NOP-tTA;tetO-ChIEF-Citrine or pOXR1-Cre mice with GIN mice that express GFP in a subset of somatostatin-expressing (SST) oriens lacunosum-moleculare (O-LM) interneurons. To silence PV interneurons, we crossed NOP-tTA;tetO-ChIEF-Citrine or pOXR1-Cre mice with a homozygous cross of PV-Cre and Ai35D, a Cre-dependent reporter mouse line that expresses archaerhodopsin-GFP (Arch-GFP).

### Stereotaxic injection of rAAV-hSyn-Flex-oChIEF-mCitrine into the MEC of pOXR1-Cre mice

To express oChIEF-mCitrine in pOXR1-Cre mice, a recombinant adeno-associated virus (rAAV, serotype 1, 4.8×10^13^ VC/ml titer) expressing FLEXed oChIEF-mCitrine (Addgene #50973) was package by Vector Biolabs (Malvern, PA). Mice were initially anesthetized via intraperitoneal injection of ketamine (100 mg/kg IP) and xylazine (2.5 mg/kg IP). Anesthesia was maintained with O_2_ supplemented with 1.0% isoflurane. For injections into the right MEC, an incision was made in the skin along the midsagittal suture, and a small hole was drilled in the skull overlying the MEC. An aluminosilicate glass pipette containing rAAV-hSyn-Flex-oChIEF-mCitrine was lowered to the level of the MEC and infused at a rate of 100 nl/min using a software driven injectomate (Neurostar, Sindelfingen, Germany). In total, 4 x 100 nl injections into the right MEC were made at AP = 300 μM rostral to the transverse sinus, ML = 3.2, and DV 4.0 to 3.55 mm. 21-28 days post viral injection, 61-105 day old mice were sacrificed for experimentation.

### Preparation of hippocampal slices

Brain slices were obtained by methods previously described (Bell et al., 2011). Mice were anaesthetized with an intraperitoneal injection of ketamine (200 mg/kg) and xylazine (20 mg/kg). Mice were transcardially perfused with ice cold saline (consisting of (in mM): Sucrose 230, KCl 2.5, CaCl 2, MgCl_2_ 6, NaHPO_4_ 1, NaHCO_3_ 25, glucose 25) and sacrificed by decapitation. The brain was removed, hemi-sected, and horizontal slices containing the mid temporal hippocampus were cut at 350 μm on a Leica VT1200 (Leica Microsystems, Buffalo Grove, IL). Sections were incubated in a holding chamber kept at 34°C. The holding chamber solution consisted of normal saline (in mM): NaCl 125, KCl 3.0, CaCl 1.2, MgCl_2_ 1.2, NaHPO_4_ 1.2, NaHCO_3_ 25, glucose 25 bubbled with 95% O_2_ / 5% CO_2_. Recordings were performed at 32-35°C.

### Light-evoked release of glutamate from MEC axon terminals and light-evoked silencing of CA1 interneuron subtype populations

MEC terminals expressing oChIEF-mCitrine were stimulated by blue light and interneurons expressing Arch-GFP were hyperpolarized by orange light. Both light paths were transmitted through the epi-illumination light path of an Olympus BX51WI microscope and a 20x water immersion objective (0.95 NA). Blue light flashes (0.1 ms in duration) and orange light pulses (3 s in duration) were generated from light-emitting diodes (LEDs) (UHP-microscope-LED-460 or UHP-T-LED-White filtered by an HQ 605/50x excitation filter, respectively, Prizmatix Modiin-Ilite, Givat Shmuel, Israel). Blue or orange light exiting the LEDs were reflected or passed through a dichroic mirror (515dcxru, Chroma Technology, Bellows Falls, VT, USA) using an optiblock beam combiner (Prizmatix) and were focused into the epi-illumination light path of an Olympus BX51WI microscope and back aperture of a 20x water immersion objective (0.95 NA) by a dichroic mirror (700dcxxr, Chroma Technology, Bellows Falls, VT, USA) in the filter turret. To release glutamate from alvear terminals, three protocols were used. To assess alvear synaptic function, 10 or 20 flashes of light (0.1 ms duration at 50 ms intervals) were delivered through the objective and focused onto CA1 SO and adjacent neocortex. CA1 SR and SLM were not exposed to light. To determine alvear excitability of CA1 PCs and interneurons to theta burst stimulation, 0.1 ms flashes of blue light were delivered to the SO at either 5 Hz (10 bursts separated by 200 ms intervals with each burst consisting of 5 pulses of light at 50 Hz) or 10 Hz (10 bursts separated by 100 ms intervals with each burst consisting of 5 pulses of light at 100 Hz).

### Electrophysiological measurements

Whole cell patch clamp recordings from hippocampal CA1 interneurons were performed using patch pipettes (2 - 4 MΩ) pulled from borosilicate glass (8250 1.65/1.0 mm) on a Narishige PC-10 pipette puller filled with (in mM): KMeSO_4_ 140, NaCl 8, MgATP 2, NaGTP 0.1, HEPES 10, and biocytin 0.1% or CsMeSO_3_ 120, NaCl 8, MgATP 2, NaGTP 0.1, HEPES 10, BAPTAK_4_ 10, QX 314 chloride 10, biocytin 0.1%. Membrane potentials and/or currents were measured with a Model 2400 patch clamp amplifier (A-M Systems, Port Angeles, WA) and converted into a digital signal by a PCI-6040E A/D board (National instruments, Austin, TX). WCP Strathclyde Software was used to store and analyze membrane potential and current responses on a PC computer (courtesy of Dr. J Dempster, Strathclyde University, Glasgow, Scotland). Calculated junction potentials (9.4 and 10 mV, respectively) were not compensated for in the analysis. Further analysis was performed with OriginPro 2018 (OriginLab Corp., Northampton, MA, USA) and Graphpad Prism (San Diego, CA).

### Morphological reconstruction of interneurons

Following electrophysiological recordings, slices were fixed in 4% paraformaldehyde (Boston BioProducts) and incubated with streptavidin Alexa Fluor 633 (ThermoFisher Scientific) in phosphate buffered saline (PBS) with Triton-X 100 as previously described (Bell et al., 2011). Processed slices were then reconstructed using a Zeiss LSM 710 confocal microscope (Carl Zeiss, Jena, Germany). Alexa Fluor 633 was excited with the 633 nm line of a HeNe 5 mW laser and cells were visualized using a 20x dry lens (0.8 N.A., voxel dimensions 0.2 x 0.2 x 1.1 µm).

### Statistics and data analysis

Data were analyzed using WCP software and OriginPro 2018 for the electrophysiological measurements. Statistics were performed using GraphPad Prism (San Diego, CA). Statistical significances for changes in the amplitude of excitatory postsynaptic currents (EPSCs) during stimulation trains were determined using a repeated measures one-way ANOVA with Bonferroni post hoc tests. Statistical significances of the effects of antagonists on EPSC and inhibitory postsynaptic current (IPSC) normalized amplitudes were determined by one sample t-tests. Comparisons between the excitability of interneuron subtypes to alvear input stimulation were assessed by two-way ANOVA with Bonferroni post hoc tests. Differences were determined to be statistically significant for *P* values less than 0.05. All data was reported as the mean, standard error of the mean (SEM). Asterisks were as follows *** *P* < 0.001, ** *P* < 0.01, * *P* < 0.05.

### Chemicals

All chemicals were purchased from VWR unless otherwise indicated. QX314 chloride and 4-aminopyridine (4-AP) were from Sigma-Aldrich, (St. Louis, MO). Tetrodotoxin citrate (TTX), bicuculline methochloride, 2,3-Dioxo-6-nitro-1,2,3,4-tetrahydrobenzo[f]quinoxaline-7-sulfonamide disodium salt (NBQX), DL-2-Amino-5-phosphono pentanoic acid (APV) were purchased from Hello Bio (Princeton, NJ). Biocytin (B-1592) and streptavidin Alexa 633 were purchased from ThermoFisher Scientific.

## Results

Alvear pathway axons travel through the alveus and adjacent CA1 SO to form varicosities that presumably form synaptic connections with CA1 PC basal dendrites and inhibitory interneuron processes located in the deep layers of hippocampal CA1 (Deller T *et al*. 1996). However, little is known about the physiological impact of the alvear pathway. Therefore, we investigated the effect of alvear pathway synaptic inputs on both CA1 PCs and inhibitory interneurons in the deep layers of hippocampal CA1.

### MEC alvear inputs form subthreshold facilitating excitatory synaptic connections with CA1 pyramidal neurons

We first assessed the effect of alvear pathway synaptic inputs onto CA1 PCs. To do this we used two different transgenic mouse lines to express the optogenetic activator protein oChIEF (Lin JY *et al*. 2009) in MEC L3 projection neurons. The first transgenic line expressed Cre-recombinase under the control of the oxidation resistance protein 1 promoter, pOXR-1-Cre (Suh J *et al*. 2011). This mouse line expresses Cre-recombinase in L3 neurons of the MEC. In this line, we intracranially injected the MEC with an adeno-associated virus (AAV) serotype 1 that expressed oChIEF-mCitrine in a Cre-dependent manner. The second transgenic mouse line expressed the tetracycline transactivator (tTA) under the control of the neuropsin promoter (NOP-tTA) (Yasuda M and MR Mayford 2006). The NOP-tTA line expresses the tetracylcline transactivator in MEC L2 neurons as well as a smaller population of MEC L3 neurons. We expressed oChIEF-mCitrine in MEC L2 and 3 neurons by crossing the NOP-tTA line with a transgenic tTA reporter mouse line in which oChIEF-mCitrine was controlled by the tetracycline transactivator (TRE) (Cheetham CEJ *et al*. 2016). This cross (NOP-tTA;TRE-oChIEF-mCitrine) resulted in the expression of oChIEF-mCitrine in MEC L2 and 3 projection neurons. For experiments examining alvear pathway inputs onto CA1 pyramidal neurons, we used 37 pOXR-1-Cre mice (15 male and 22 female) and 12 NOP-tTA;TRE-oChIEF-mCitrine mice (7 male and 5 female). A total of 41 of 49 pyramidal neurons responded to optogenetic stimulation of the alvear pathway. Data were pooled for analysis.

We first assessed the electrophysiological properties of CA1 pyramidal neurons before examining their responses to alvear inputs (Table 1). The electrophysiological properties of CA1 pyramidal neurons consisted of resting membrane potential averages of -65.2 ± 0.9 mV with average action potential amplitudes of 85.8 ± 2.8 mV and durations of 1.8 ± 0.1 ms (N = 20). CA1 pyramidal neurons produced outwardly rectifying current-voltage relationships to hyperpolarizing current injection (Fig. 1A, B). Larger hyperpolarizing current injections produced a depolarizing sag in the membrane potential response. Suprathreshold depolarizing currents resulted in adapting trains of action potentials with small afterdepolarizations (Fig. 1A). In voltage clamp, activation of the alvear pathway with 20 Hz pulses of blue light (0.1 ms duration) resulted in facilitating EPSCs (Fig. 1C, D). EPSC amplitudes displayed a significant linear trend to larger successive EPSC amplitudes (repeated-measure ANOVA, *P* < 0.0001, n = 7). Alvear pathway synaptic inputs onto CA1 pyramidal neuron basal dendrites therefore showed a low probability of neurotransmitter release.

**Figure 1.**
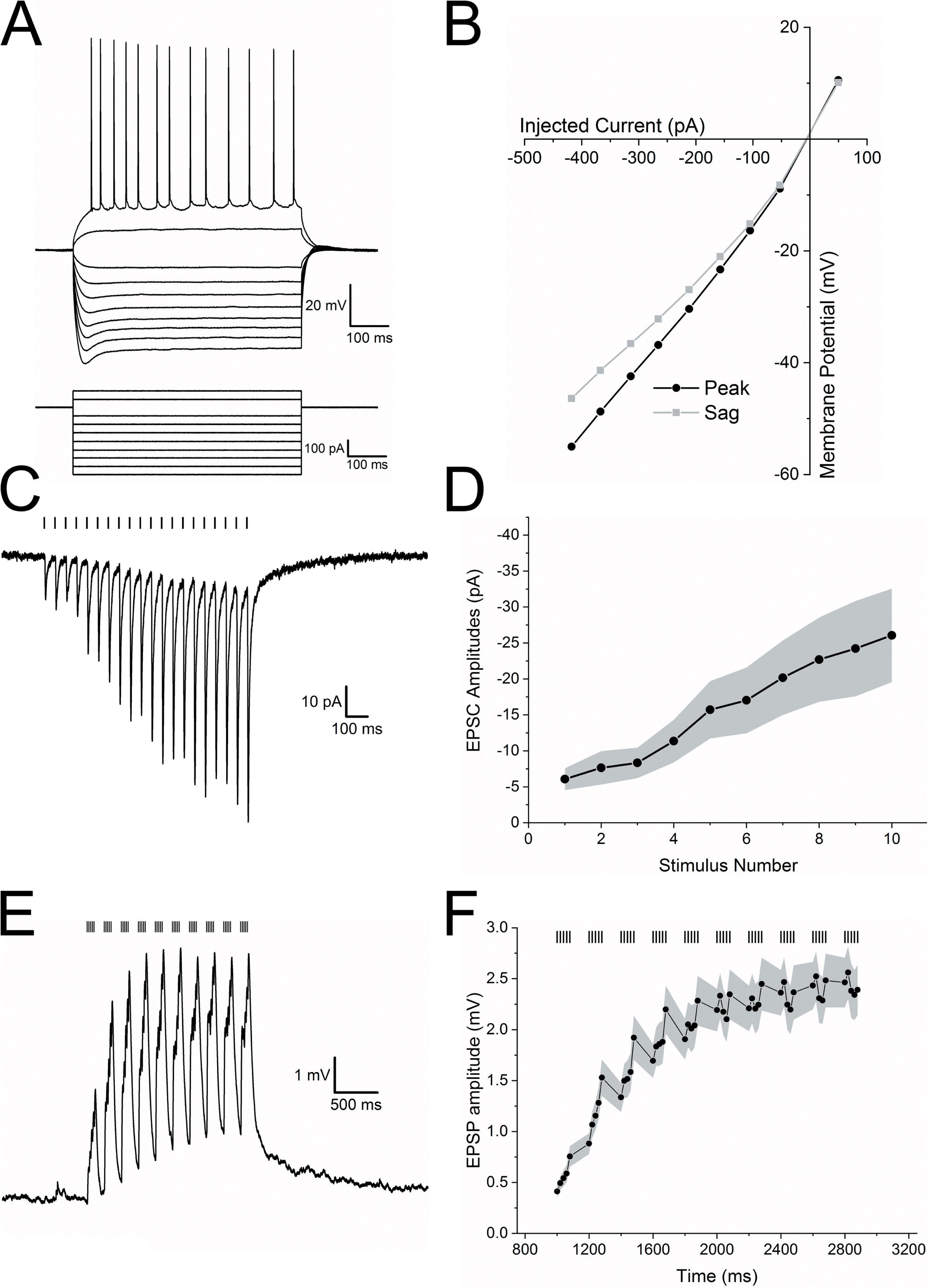
The alvear pathway forms functional synaptic connections onto CA1 pyramidal neurons. **A**. CA1 pyramidal cell (PC) membrane potential responses to a series of current injections of varying amplitudes (600 ms duration). **B**. Current-voltage relationship demonstrating slight outward rectification and depolarizing sag in the membrane potential of a CA1 PC following hyperpolarizing current injections. Black line – peak voltage response amplitude, grey line – voltage response amplitude at the end of the current pulse. **C**. Optogenetic activation of the alvear pathway produced facilitating EPSCs (20 light pulses, 0.1 ms duration, 50 ms intervals) in an individual CA1 PC. Bars above current trace indicate timing of blue light flashes. **D**. Line plot demonstrating that the normalized average EPSC amplitude in CA1 PCs significantly facilitated in response to 20 Hz stimulation. Shaded area shows SEM. **E**. Theta burst optogenetic stimulation of the alvear pathway produced subthreshold facilitating EPSPs in an individual CA1 PC. Bars above the voltage trace indicates the timing of blue light flashes. **F**. Line plot of the average EPSP amplitude in response to theta burst stimulation of the alvear pathway. EPSP amplitudes significantly facilitated demonstrating a linear trend to larger amplitudes. Shaded area indicates SEM.

**Table 1.** Electrophysiological properties of known cell types from which recordings were made in this study.

| **Cell Type** | **AP**<br>**Amplitude**<br>**(mV)** | **AP**<br>**threshold**<br>**(mV)** | **AP**<br>**duration**<br>**(ms)** | **AP mean**<br>**frequency**<br>**(Hz)** | **AP**<br>**frequency**<br>**ratio**<br>**(first/last)** | **Resting**<br>**Membrane**<br>**Potential (mV)** | **Input**<br>**Resistance**<br>**(mΩ)** | **I<sub>h</sub> Sag**<br>**Ratio** |
| --- | --- | --- | --- | --- | --- | --- | --- | --- |
| **CA1**<br>**Pyramidal**<br>**Cell (n = 20)** | 85.8<br>± 2.8 | -43.9<br>± 0.8 | 1.77<br>± 0.11 | 35.9<br>± 3.0 | 0.36<br>± 0.04 | -65.2<br>± 0.9 | 180.7<br>± 18.7 | 0.82<br>± 0.01 |
| **PV**<br>**Interneuron (n**<br>**= 58)** | 69.0<br>± 1.5 | -42.8<br>± 0.9 | 0.51<br>± 0.01 | 184.8<br>± 7.1 | 1.30<br>± 0.23 | -61.6<br>± 0.9 | 111.0<br>± 7.6 | 0.93<br>± 0.005 |
| **VIP**<br>**Interneuron**<br>**(n = 16)** | 74.3<br>± 2.4 | -40.8<br>± 0.8 | 1.01<br>± 0.03 | 64.1<br>± 8.5 | 0.42<br>± 0.04 | -59.3<br>± 1.0 | 492.7<br>± 58.6 | 0.84<br>± 0.02 |
| **GIN**<br>**Interneuron (n**<br>**= 29)** | 64.0<br>± 2.5 | -38.4<br>± 1.2 | 0.90<br>± 0.04 | 65.2<br>± 4.8 | 0.74<br>± 0.09 | -57.0<br>± 1.3 | 394.7<br>± 42.0 | 0.84<br>± 0.02 |

The activity of individual MEC L3 projection neurons *in vivo* has been poorly correlated with population theta rhythm activity (Quilichini P et al. 2010). Nevertheless, during theta rhythms the largest current sink in hippocampal CA1 occurs in the SLM and is eliminated following blockade of EC inputs (Bragin A et al. 1995; Kamondi A et al. 1998). Because the alvear pathway projects to hippocampal CA1 SLM, alvear inputs likely contribute to the sources and sinks observed during theta rhythm. Furthermore, individual grid cells located in L3 of the MEC fire action potentials correlated with their own cell’s theta rhythms (Quilichini P *et al*. 2010; Domnisoru C et al. 2013). Therefore, we examined whether theta burst activation of alvear pathway boutons in the SO of hippocampal CA1 were capable of activating CA1 pyramidal neurons. Although CA1 PCs responded by producing EPSPs, none of the 26 CA1 pyramidal neurons from which we recorded produced action potentials in response to theta burst stimulation (Fig. 1E). On average, the summating EPSPs (linear trend, repeated-measures ANOVA, *P* < 0.0001, n = 26) did not exceed 3 mV in amplitude (Fig. 1F). Therefore, although the alvear pathway appears to innervate the basal dendrites of CA1 pyramidal neurons, synaptic release from the alvear input resulted in small subthreshold EPSPs measured at the pyramidal cell soma.

### MEC alvear inputs monosynaptically excite and disynaptically inhibit CA1 pyramidal neurons

TA inputs from the MEC arise from both inhibitory and excitatory neurons (Melzer S et al. 2012). However, it is unknown whether synaptic connections of the alvear pathway onto hippocampal neurons are excitatory or inhibitory. Therefore, we measured both EPSCs and IPSCs generated by theta burst stimulation of alvear inputs onto CA1 pyramidal neurons. For these studies we used 15 pOXR1-Cre mice (5 female, 10 male) and 14 NOP-tTA mice (4 female, 10 male). Data from all mice were pooled.

Optogenetic theta burst stimulation produced summating EPSCs when CA1 pyramidal neurons were voltage clamped at -65 mV, near the reversal potential for chloride (Fig. 2A, black trace). Application of TTX (1 μM, Fig. 2A, red trace) resulted in partial inhibition of the EPSCs (Fig. 2E, repeated measures ANOVA, Bonferroni post hoc test, *P* < 0.0001, n = 3). Subsequent, application of 4-aminopyridine (100 μM, 4-AP) with TTX moderately rescued the EPSCs (Fig. 2E, repeated measures ANOVA, Bonferroni post hoc test, *P* = 0.0019, n = 3), particularly during the initial stimuli in a burst (Fig. 2A, green trace). These observations suggest that the EPSCs in CA1 pyramidal neurons resulted from monosynaptic inputs from the MEC alvear pathway (Petreanu L et al. 2007).

**Figure 2.**
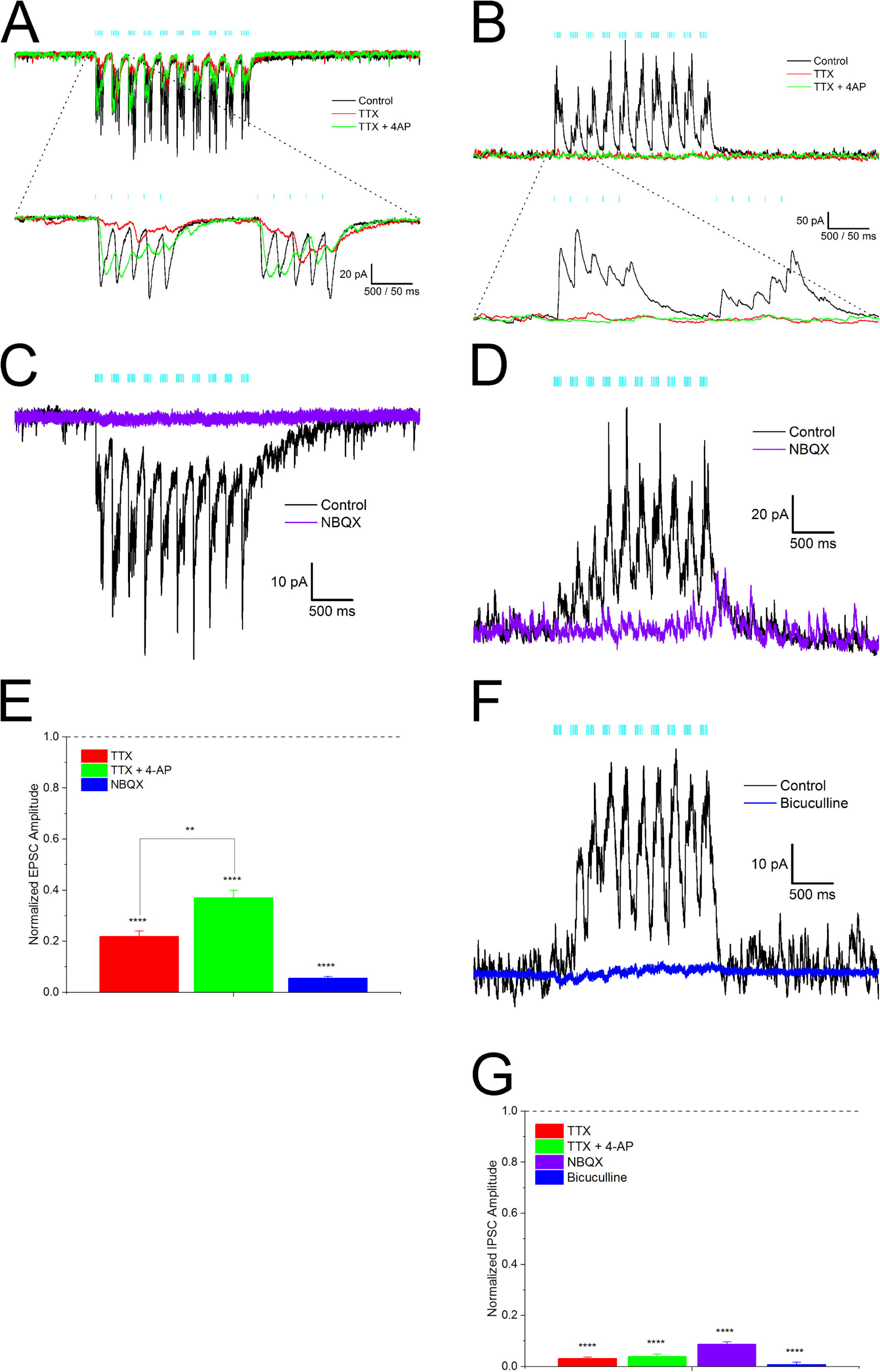
Alvear pathway forms monosynaptic excitatory and disynaptic inhibitory connections onto CA1 pyramidal neurons. **A**. Theta burst optogenetic stimulation consisting of 10 bursts of blue light pulses (5 pulses, 0.1 ms duration, 50 ms intervals) separated by 200 ms resulted in EPSCs (black trace) in a voltage-clamped PC (Vh = -65 mV). EPSCs were partially inhibited by TTX (1 μM, red trace). Subsequent co-application of 4-AP (100 μM, green trace) partially rescued EPSC amplitudes. Bottom traces – expansion of the first two optogenetic bursts in the theta burst sequence. **B.** Same pyramidal neuron in A voltage clamped at +15 mV. Theta burst stimulation produced outward IPSCs (black trace) that were completely inhibited by TTX (1 μM, red trace). Subsequent addition of 4-AP (100 μM, green trace) did not rescue the TTX blockade. Bottom traces – expansion of the first two optogenetic bursts in the theta burst sequence. **C.** NBQX (30 μM, purple trace) completely blocked theta burst driven EPSCs (black trace) in a CA1 pyramidal neuron (Vh = -65 mV). **D.** NBQX (30 μM) completely inhibited the theta burst driven outward IPSCs in a CA1 pyramidal neuron (Vh = +15 mV). **E.** Summary bar plot of the effect of TTX, TTX + 4-AP, and NBQX on the summated EPSC amplitudes. Burst amplitudes were averaged and normalized to control amplitudes. **F.** Bicuculline (25 μM, blue trace) completely abolished the outward IPSC (black trace) in a pyramidal neuron. **G.** Summary bar plot of the effect of TTX, TTX + 4-AP, NBQX, and bicuculline on averaged summated IPSC amplitudes. Amplitudes were normalized to control.

To record inhibition produced by MEC alvear stimulation, the membrane potential was held at approximately +15 mV near the reversal potential for ionotropic glutamate receptors. Optogenetic theta burst stimulation at +15 mV resulted in outward IPSCs (Fig. 2B, black trace) that were completely blocked by TTX (1 μM, Fig. 2B and G, red, repeated measures ANOVA, Bonferroni post hoc test, *P* < 0.0001, n = 3). However, unlike alvear EPSCs, the IPSCs were not rescued by subsequent addition of 4-AP (100 μM, Fig. 2B and G, green). These data suggest that the MEC alvear pathway can inhibit CA1 pyramidal neurons indirectly through the excitation of inhibitory interneurons.

We next examined the receptor subtypes mediating postsynaptic responses in CA1 PCs. Application of the AMPA receptor antagonist NBQX (30 μM) completely inhibited EPSCs (Fig. 2C and E, purple, one sample t-test, *P* < 0.0001, n = 3) and IPSCs (Fig. 2D and G, purple, one sample t-test, *P* < 0.0001, n = 3) elicited by the optogenetic stimulation of alvear inputs. Optogenetic theta burst driven IPSCs could also be inhibited by the application of the GABA_A_ receptor antagonist bicuculline at 25 μM (Fig. 2F and G, blue, one sample t-test, *P* < 0.0001, n = 10). Therefore, these data suggest that AMPA receptors mediated both the monosynaptic excitatory inputs and the indirect activation of inhibitory inputs onto CA1 PCs by the alvear pathway. Interneurons excited by alvear inputs then inhibit postsynaptic CA1 PCs via the activation of GABA_A_ receptors.

MEC alvear inputs excite horizontally-oriented interneurons in stratum oriens of hippocampal CA1

Although activation of alvear inputs do not appear to be potent enough to elicit action potentials in CA1 PCs, the same activation does elicit inhibitory synaptic responses in CA1 PCs (Fig. 2). Therefore, we next examined the effect of optogenetic stimulation of alvear inputs on inhibitory interneuron activity in the SO of hippocampal CA1. In total, we recorded from 39 interneurons with somas located in hippocampal CA1 SO. We included biocytin in the patch pipette to aid in the identification of these interneurons. Out of 39 interneurons, 11 did not respond. Of the remaining 28 interneurons, 15 could not be reconstructed, and the remaining 13 had soma and dendrites confined to the SO (eg. Fig. 3D) demonstrating that these interneurons could only be activated by axon terminals in the SO and not the SLM.

**Figure 3.**
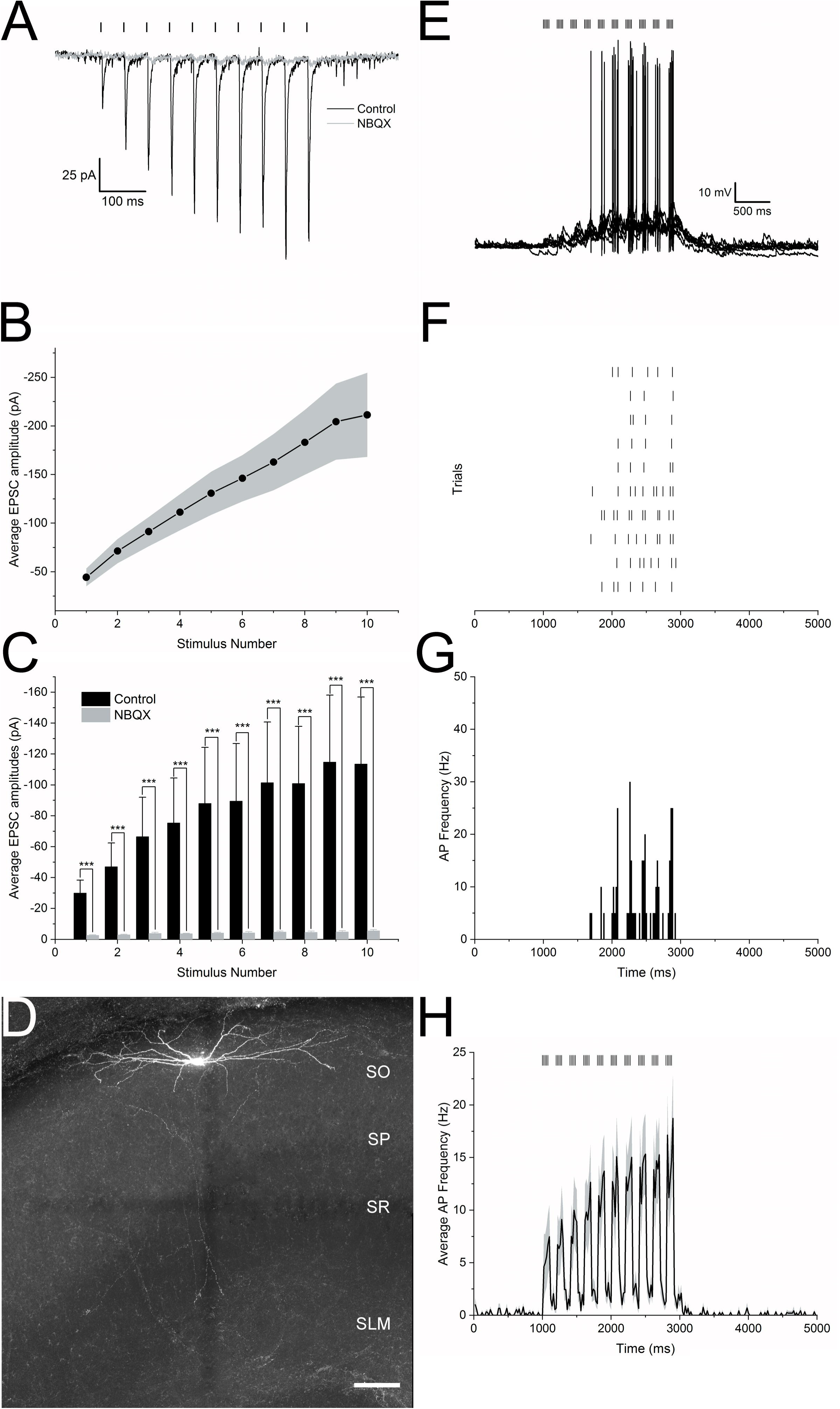
Alvear inputs excite CA1 inhibitory interneurons in the alveus, stratum oriens, and stratum pyramidale. **A**. Optogenetic activation of MEC alvear inputs (10 x 0.1 ms blue light pulses at 20 Hz, indicated by bars above traces) produced facilitating EPSCs (black trace) in a CA1 stratum oriens interneuron. Application of the AMPA receptor antagonist NBQX (30 μM) inhibited the EPSCs (grey trace). **B**. Line plot demonstrating a consistent facilitation of alvear EPSCs (10 pulses at 20 Hz) in CA1 inhibitory interneurons. Grey shading represents the SEM. **C.** Bar plot demonstrating that alvear EPSCs (10 pulses at 20 Hz) were reliably inhibited by the AMPA receptor antagonist NBQX. **D**. Confocal image of a horizontal CA1 inhibitory interneuron that was excited by optogenetic activation of alvear inputs and has its soma and dendrites confined to the SO. Scale bar 100 μm. **E**. 5 Hz optogenetic theta burst stimulation of alvear inputs produced suprathreshold responses in a CA1 SO interneuron. Bars above 10 superimposed traces indicate timing of light flashes. **F**. Raster plot of the timing of action potentials occurring during each trial taken from the sweeps of E. **G**. Peristimulus time histogram showing the mean action potential firing frequency that occurred during 20 ms bins in the interneuron of F and G. **H**. Peristimulus time histogram of the mean action potential firing frequency (20 ms bins) averaged from 22 different CA1 alvear/stratum oriens/stratum pyramidale interneurons in response to 5 Hz optogenetic stimulation of the alvear pathway. Grey shading indicates the SEM.

We first examined the strength of alvear inputs by optogenetically stimulating their afferents (10 blue light pulses, 0.1 ms duration, 50 ms intervals) and assessing short-term synaptic dynamics. All interneurons responded with facilitating EPSCs (Fig. 3A, black trace). The EPSC facilitation showed a significant linear trend to larger amplitudes with subsequent stimuli (Fig. 3B, repeated-measures ANOVA, linear trend, *P* < 0.0001, n = 8). EPSCs in all interneurons were blocked by the glutamatergic AMPA receptor antagonist NBQX at 30 μM (Fig. 3A, C, B grey trace/bar, two-way ANOVA, Bonferroni post hoc test, *P* < 0.0001, n = 10). Therefore, the alvear inputs appear to have a low probability of release and activate AMPA receptors on horizontally oriented interneurons (Fig. 3D) in hippocampal CA1.

We next examined the excitability of CA1 SO neurons to 5 Hz theta burst optogenetic stimulation. Of the 28 interneurons that responded to alvear stimulation, 17 interneurons fired action potentials during theta burst stimulation (Fig. 3E-G). Of those 17 activated interneurons, seven had sufficient biocytin reconstruction to show soma and dendrites confined to stratum oriens with two having axonal fills consistent with perisomatic interneurons and one with bistratified-like axonal arborization. Averaging the peristimulus time histogram data among all responding SO interneurons demonstrated that the average frequency of action potentials increased with subsequent optogenetic bursts (Fig. 3H). Therefore, these data demonstrated that alvear inputs could excite inhibitory interneurons in hippocampal CA1 and produce feedforward inhibition of CA1 pyramidal neurons.

### Parvalbumin-expressing hippocampal CA1 interneurons can be activated by alvear inputs

TA inputs from the entorhinal cortex innervate hippocampal CA1 in the SLM. Stimulation of the TA has been shown to activate interneurons that in turn activate both GABA_A_ and GABA_B_ receptors on CA1 PCs (Empson RM and U Heinemann 1995, 1995; Dvorak-Carbone H and EM Schuman 1999; McQuiston AR 2011; Milstein AD *et al*. 2015). However, stimulation of axons in the SLM failed to activate parvalbumin-expressing (PV) interneurons (Milstein AD *et al*. 2015). These latter studies suggest that the entorhinal cortex may play a limited role in inhibiting CA1 PC output and may not contribute to behaviorally-relevant rhythmicity as PV interneurons play major roles in both these processes (Pelkey KA et al. 2017). However, alvear pathway terminals in CA1 are located in a position where they could activate PV interneurons. The next set of experiments examined whether PV interneurons in CA1 responded to alvear pathway stimulation.

To target PV interneurons, we crossed NOP-tTA;oChIEF-mCitrine or pOXR-1-Cre mice to transgenic mice expressing tdTomato under the control of the parvalbumin promoter (Kaiser T *et al*. 2016) or homozygous crosses of PV-Cre (Hippenmeyer S *et al*. 2005) and Ai14 (Madisen L *et al*. 2010). Recordings from tdTomato-positive (PV) interneurons displayed fast non-adapting action potentials when depolarized by current injection. These interneurons also produced slightly outwardly rectifying hyperpolarizations with little depolarizing sag in response to hyperpolarizing current injection (Fig. 4A). The electrophysiological properties of these interneurons consisted of resting membrane potential averages of -61.2 ± 0.8 mV, input resistances of 111.0 ± 7.7 MΩ, 69.0 ± 1.5 mV action potential amplitudes and 0.51 ± 0.01 ms action potential durations (Table 1, n = 58). Post hoc anatomical reconstruction of biocytin-filled PV interneurons showed that 29 cells had perisomatic axonal arborizations (Fig. 4Aiii), five had bistratified morphology (Fig. 4Aiv) and 24 did not have sufficient axonal fill to determine anatomical subtype.

**Figure 4.**
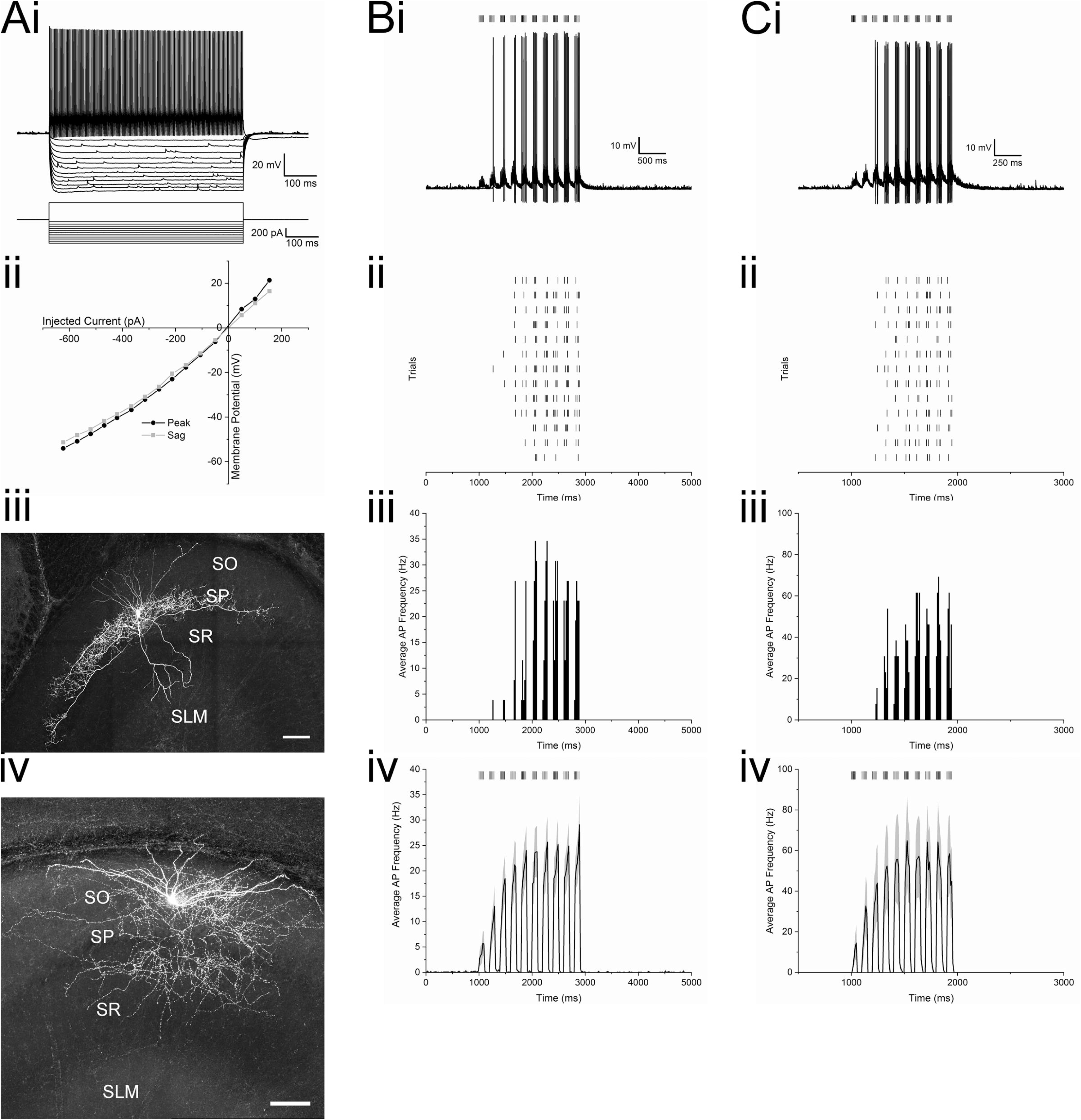
Alvear inputs excite hippocampal CA1 parvalbumin-expressing interneurons. **Ai.** Membrane potential responses of a PV interneuron (top) to steps of current injection (bottom). **Aii.** Line plot illustrating the membrane potential amplitude responses to current injections (black line - maximum voltage response (peak), grey line – membrane potential amplitude at end of response (sag)). **Aiii.** Confocal image of a PV interneuron with perisomatic axonal arborization that responded to alvear stimulation. Scale bar 100 μm. **Aiv.** Confocal image of a PV interneuron with bistratified axonal arborization that responded to alvear stimulation. Scale bar 100 μm. **Bi.** 13 superimposed traces of the response of a PV interneuron to 5 Hz optogenetic theta burst stimulation. **Bii**. Raster plot of the timing of action potentials in response to optogenetic stimulation shown in Bi. **Biii.** Peristimulus time histogram demonstrating the mean action potential firing frequency (20 ms bins) in response to optogenetic theta burst stimulation shown in Bi and Bii. **Biv.** Peristimulus time histogram of the mean action potential firing frequency (20 ms bins) averaged from 29 PV interneurons that responded with action potentials to alvear pathway 5 Hz optogenetic theta burst stimulation. Grey shading indicates SEM. **Ci**.13 superimposed traces of membrane potential responses of the same PV interneuron in B to 10 Hz optogenetic theta burst stimulation. **Cii.** Raster plot illustrating the timing of action potentials in response to 10 Hz optogenetic theta burst stimulation from the interneuron shown in Ci. **Ciii**. Peristimulus time histogram illustrating the mean action potential firing frequency (10 ms bins) that occurred in the interneuron shown in Ci and Cii to 10 Hz optogenetic theta burst stimulation. **Civ**. Averaged peristimulus time histogram of the mean action potential firing frequency taken from 29 PV interneurons in response to 10 Hz optogenetic theta burst stimulation. Grey shading indicates the SEM.

We next examined whether action potentials could be produced in PV interneurons by optogenetic theta burst activation of alvear pathway inputs. Both 5 Hz and 10 Hz optogenetic theta burst stimulation of the alvear pathway resulted in the production of action potentials in 29 interneurons. Of the interneurons that produced suprathreshold responses, 18 were perisomatic (Fig. 4Aiii), one was bistratified (Fig. 4Aiv), and 10 had insufficient biocytin fills for anatomical identification. In each responsive interneuron, 5 Hz or 10 Hz optogenetic theta burst stimulation reliably produced action potentials during each trial (Fig. 4Ai-iii, Bi-iii). When all action potential responsive interneurons were averaged and peristimulus time histograms constructed, 5 Hz optogenetic theta burst stimulation resulted in peak burst frequencies near 25 Hz whereas 10 Hz theta burst produced burst frequencies of near 60 Hz (Fig. 4Aiv, Biv). Therefore, unlike the perforant path (Milstein AD *et al*. 2015), both low frequency (5 Hz) and high frequency (10 Hz) theta burst activation of the entorhinal cortical alvear pathway was capable of activating both perisomatic and bistratified PV interneurons in hippocampal CA1.

### VIP-expressing CA1 hippocampal interneurons can be activated by alvear inputs

In addition to PV-expressing neurons, a subset of VIP-expressing interneurons constitute another class of perisomatic projecting interneurons onto CA1 PCs (Klausberger T and P Somogyi 2008; Pelkey KA *et al*. 2017). In order to target VIP-expressing cells for the next set of experiments, we crossed either NOP-tTA;oChIEF-mCitrine or pOXR-1-Cre mice with homozygous crosses of a VIP-Cre driver mouse line (Taniguchi H *et al*. 2011) and a tdTomato Cre-dependent reporter mouse line (VIP-Cre;ai14) (Madisen L *et al*. 2010). pOXR-1-Cre;VIP-Cre;ai14 crosses were injected with AAV-hSyn-Flex-oChIEFmCitrine into the MEC. tdTomato-expressing VIP interneurons were targeted for whole cell patch clamp recordings, and electrophysiological responses to optogenetic stimulation were measured from recorded neurons.

VIP-expressing interneurons had varying electrophysiological properties. Perisomatic VIP-expressing interneurons had regular spiking firing patterns with adapting action potential frequencies (Fig. 5Ai) (Bell LA et al. 2015). Averaged electrophysiological properties of these interneurons were defined by -59.3 ± 0.1 mV resting membrane potentials, 492.7 ±58.6 MΩ input resistances, 74.3 ± 2.4 mV action potential amplitudes, and 1.01 ± 0.03 ms action potential durations. When we delivered optogenetic theta burst stimulation to 37 VIP-expressing interneurons, 16 VIP interneurons responded by producing action potentials, (Fig. 5B and C), 12 responded with subthreshold excitatory postsynaptic potentials, and nine displayed no response. Of the 16 interneurons that produced action potentials, four had perisomatic axonal arborizations, three had non-perisomatic morphology, and nine could not be identified. Of the 10 interneurons that responded with subthreshold EPSPs, two had perisomatic axonal arborizations whereas the other 10 could not be anatomically identified.

**Figure 5.**
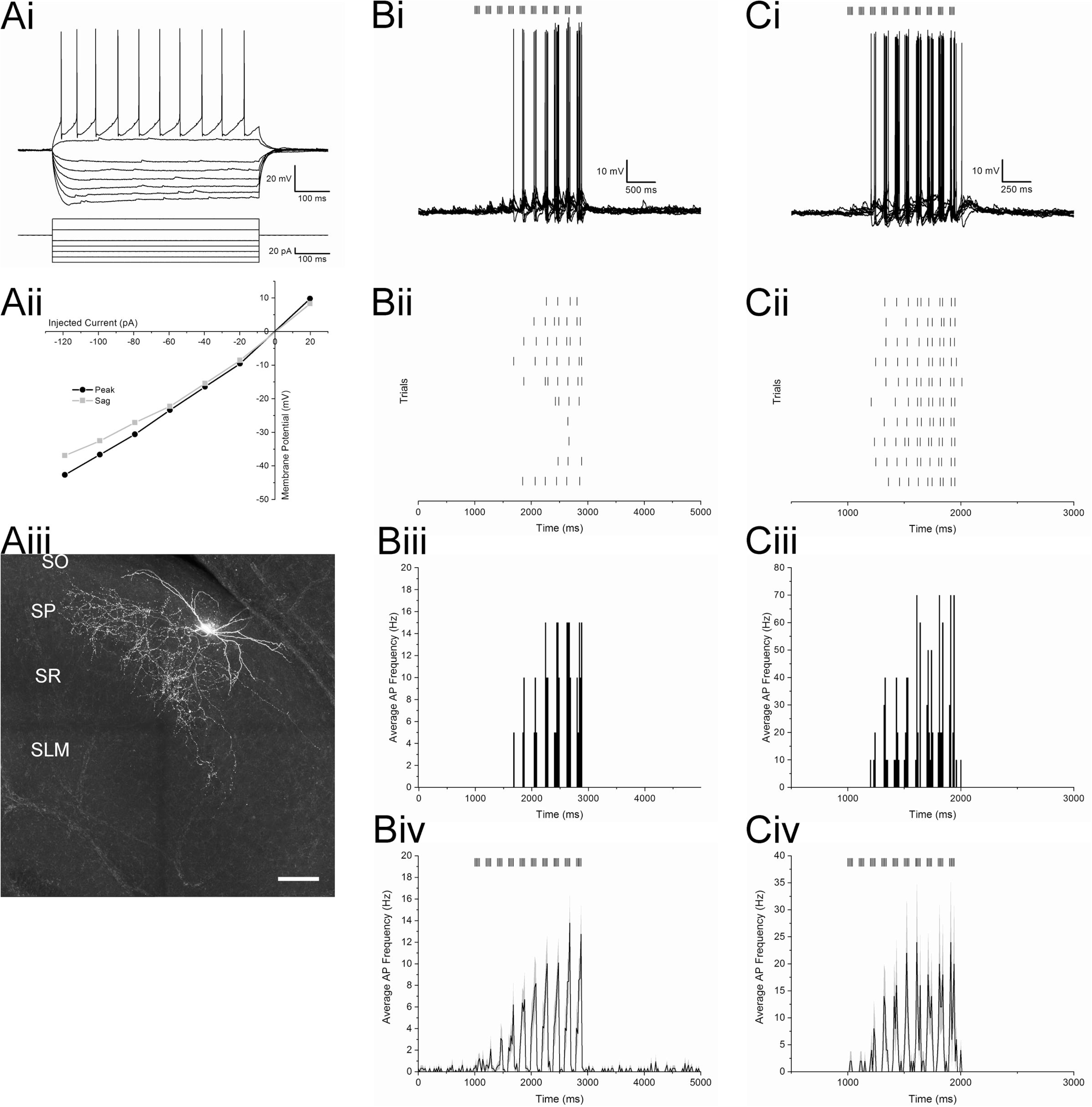
Alvear inputs excite VIP-expressing interneurons in stratum oriens of hippocampal CA1. **Ai.** Membrane potential response of a VIP interneuron to depolarizing and hyperpolarizing current injections showing slight adaptation. **Aii**. Line plot illustrating the membrane potential amplitude responses to current injections (black line - maximum voltage response (peak), grey line – membrane potential amplitude at end of response (sag)). **Aiii**. Confocal reconstruction of a VIP perisomatic interneuron that responded to alvear input stimulation. Scale bar 100 μm. **Bi**. 10 superimposed membrane potential traces of a VIP interneuron in response to 5 Hz optogenetic theta burst stimulation. Bars above the traces indicate the timing of the blue light pulses. **Bii**. Raster plot of the timing of action potentials occurring during each of the 10 individual trials of the interneuron shown in Bi. **Biii**. Peristimulus time histogram of the mean action potential firing frequency (20 ms bins) occurring during the 10 trials from the interneuron shown in Bi and Bii. **Biv**. Averaged peristimulus time histogram of the mean action potential firing frequency taken from 14 VIP interneurons that produced action potentials to 5 Hz optogenetic theta burst stimulation. **Ci**. 10 superimposed membrane potential traces of the same VIP interneuron in Bi in response to 10 Hz optogenetic theta burst stimulation. Bars above the traces indicate the timing of the blue light pulses. **Cii**. Raster plot of the timing of action potentials that occurred during the 10 trials of the neuron shown in Ci. **Ciii**. Peristimulus time histogram of the mean action potential firing frequency (10 ms bins) that occurred during the 10 trials from the neuron shown in Ci and Cii. **Civ**. Averaged peristimulus time histogram of the mean action potential frequency (10 ms bins) averaged from 14 VIP interneurons that produced action potentials to 10 Hz optogenetic theta burst stimulation.

An example of a VIP interneuron that responded to both 5 and 10 Hz optogenetic theta burst stimulation is illustrated in Figure 5 Bi and Ci, respectively. Theta burst optogenetic stimulation resulted in burst firing synchronized to the optogenetic stimulation (Fig. 5Bii, Cii). The average frequency of action potential firing is illustrated in the peristimulus time histograms (Fig. 5Biii, Ciii). When the action potential frequency of the 16 responding interneurons were averaged, the first two optogenetic bursts produced low frequencies of action potentials. The average frequency of action potentials during bursts continued to increase with subsequent optogenetic bursts peaking near 13 Hz for 5 Hz stimulation and 23 Hz for 10 Hz stimulation (Fig. 5Biv, Civ). Therefore, like PV-expressing interneurons, VIP interneurons were excited with theta burst optogenetic stimulation of alvear inputs; however, these interneurons responded with lower frequency of action potentials.

### A small population of presumptive oriens lacunosum-moleculare interneurons are activated by alvear inputs

We next examined the effect of optogenetic activation of the alvear pathway on the excitability of a subpopulation of OL-M interneurons in hippocampal CA1, which innervate the distal apical dendritic domain of CA1 PCs. Transgenic GFP interneuron mice (GIN mice) used in these studies express GFP in a subpopulation of somatostatin interneurons with axon arborizations in the stratum lacunosum-moleculare in hippocampal CA1 and CA3 (Oliva AA, Jr. *et al*. 2000). We crossed these GIN mice with either NOP-tTA;oChIEF-mCitrine or pOXR-1-Cre mice to target OL-M cells. The alvear pathway was optogenetically stimulated to assess responses in GIN cells.

The electrical properties of GIN interneurons in hippocampal slices had regular or burst spiking with adapting action potential firing patterns when depolarized by positive current steps (Fig. 6Ai). When these interneurons were hyperpolarized by negative current steps, the cell responded with outward rectification and a depolarizing sag in the hyperpolarizing membrane potential response (Fig. 6Aii). The electrophysiological properties of these interneurons consisted of resting membrane potential averages of -57.0 ± 1.3 mV, input resistances of 394.7 ± 42.0 MΩ, 64.0 ± 2.5 mV action potential amplitudes and 0.90 ± 0.04 ms action potential durations. Optogenetic theta burst stimulation of the alvear pathway induced a small subpopulation of these interneurons to fire action potentials. The GIN interneuron illustrated in Figure 6B and C responded with action potentials to both 5 and 10 Hz optogenetic theta burst stimulation (Fig. 6Bi and Ci, respectively). During each trial, this cell produced action potentials sporadically with less than 10 action potentials per sweep (Fig. 6Bii, Cii). Peristimulus time histograms showed that the mean action potential frequencies for this GIN interneuron were below 25 Hz for 5 Hz theta burst stimulation (20 ms bins, Fig. 6Biii) and 50 Hz for 10 Hz theta burst stimulation (10 ms bins, Fig. 6Ciii). For seven interneurons that responded with action potentials to 5 Hz optogenetic theta burst stimulation, the average action potential frequencies were below 15 Hz (Fig. 6Biv). Therefore, although some GIN interneurons could be excited by optogenetic theta burst stimulation, they typically responded sporadically with low frequencies of action potentials.

**Figure 6.**
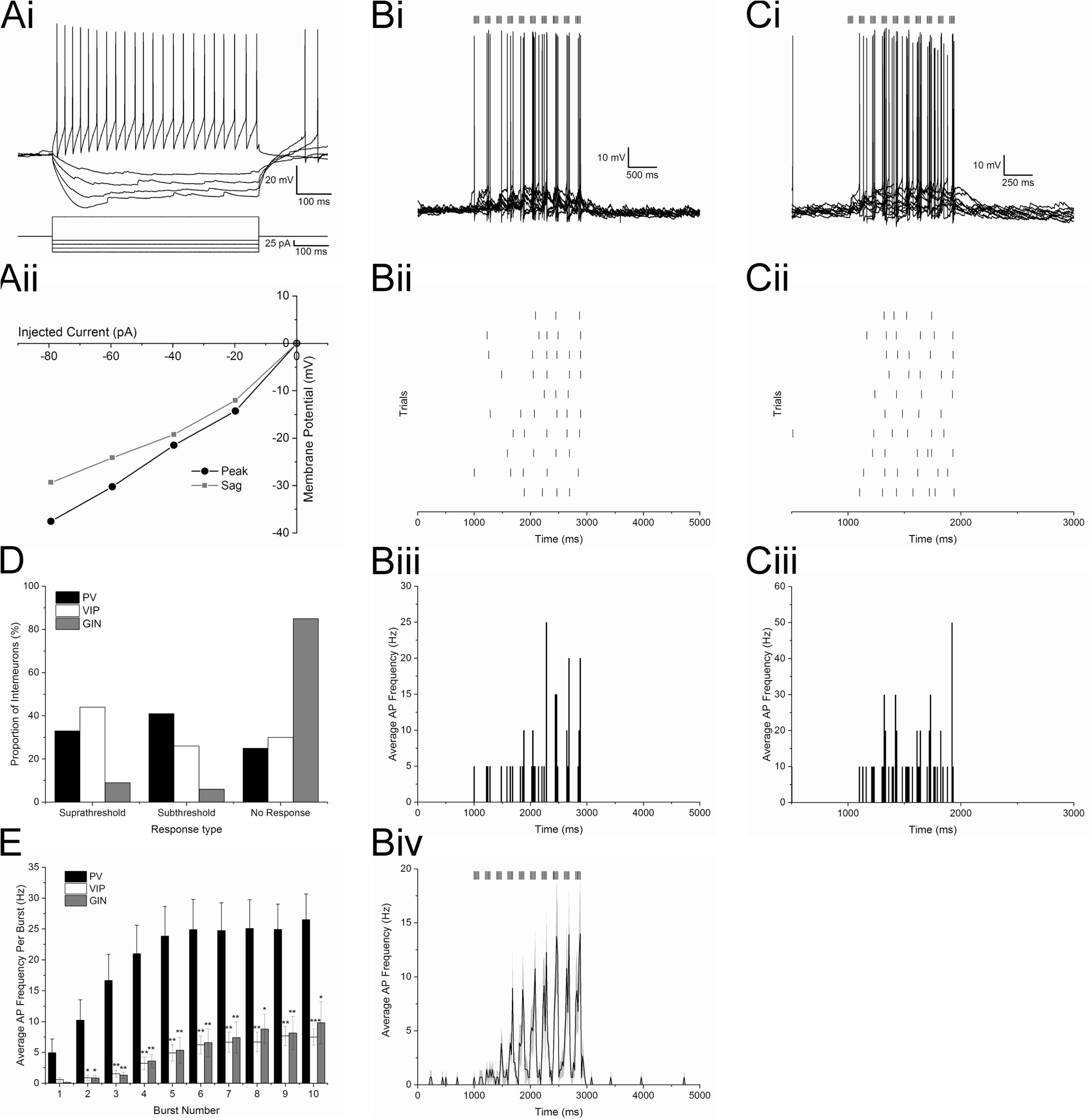
A small subset of oriens lacunosum-moleculare interneurons in hippocampal CA1 are activated by alvear inputs from the entorhinal cortex. **Ai.** Membrane potential responses of a GIN interneuron to depolarizing and hyperpolarizing current responses. **Aii**. Line plot illustrating membrane potential amplitude responses to a series of current injection steps (black line - maximum voltage response (peak), grey line – membrane potential amplitude at end of response (sag)). **Bi**. 10 superimposed traces of the response of a GIN interneuron to 5 Hz optogenetic theta burst stimulation of the alvear pathway. Vertical lines above the traces indicate the timing of the blue light flashes. **Bii**. Raster plot of the timing of action potentials in the GIN interneuron shown in Bi to 5 Hz optogenetic theta burst stimulation of the alvear pathway. **Biii**. Peristimulus time histogram (20 ms bins) illustrating the mean action potential firing frequency of the GIN interneuron shown in Bi and ii. **Biv.** Average peristimulus time histogram (20 ms bins) taken from 7 GIN neurons that responded with action potentials to 5 Hz optogenetic theta burst stimulation. **Ci**. 10 superimposed membrane potential responses, from the GIN interneuron illustrated in Bi-iii, to 10 Hz optogenetic theta burst stimulation. Vertical bars above the traces indicate the timing of blue light stimuli during each trial. **Cii**. Raster plot illustrating the timing of action potentials for each trial from the overlapping traces shown in Ci. **Ciii**. Peristimulus time histogram (10 ms bins) showing the mean action potential firing frequency of the GIN interneuron of Ci and Cii. **D**. Proportion of PV (black), VIP (white) interneurons and GIN (grey) that generated suprathreshold (left), subthreshold (middle), or no response (right) in response to 5 Hz optogenetic theta burst stimulation of the alvear pathway in hippocampal CA1. **E**. Bar graph illustrating the averaged action potential firing frequency of PV (black, n = 29), VIP (white, n = 14), or GIN (grey, n= 7) interneurons during each burst of a 5 Hz optogenetic theta burst stimulation (* *P* < 0.05, ** *P* < 0.01, *** *P* < 0.001).

In summary, the responsiveness of PV, VIP, and GIN interneurons to alvear pathway stimulation varied across subtypes. GIN neurons showed considerably less responsiveness to alvear input. Specifically, 85% of GIN neurons did not respond to alvear stimulation compared to 25% of PV and 30% of VIP interneurons that were non responsive (Fig. 6D). Of the interneurons that fired action potentials, PV interneurons produced a significantly higher frequency of action potentials during each burst of a 5 Hz optogenetic theta burst stimulation (Fig. 6E, two-way ANOVA, *P* < 0.0001, Bonferroni post hoc tests). Therefore, PV interneurons appear to be more excitable to MEC alvear inputs compared to interneurons that express VIP or somatostatin.

### Suppression of PV interneuron activity inhibits alvear-driven inhibitory postsynaptic currents in hippocampal CA1 pyramidal neurons

Hippocampal CA1 PV interneurons appear to be largely unaffected by TA inputs located in the SLM of CA1 (Milstein AD *et al*. 2015). However, both perisomatic and bistratified PV interneurons located in deep layers of hippocampal CA1 can be excited to produce action potentials by stimulation of the alvear pathway (Fig. 4). Therefore, we examined the contribution of PV interneurons to the multi-synaptic IPSCs measured in hippocampal CA1 pyramidal neurons during theta burst stimulation of alvear inputs.

To do this we crossed pOXR-1-Cre mice or NOP-tTA;oChIEF-mCitrine mice with mice that expressed the inhibitory optogenetic protein Arch-GFP in PV interneurons (PV-Cre;Ai35) (Hippenmeyer S *et al*. 2005; Madisen L *et al*. 2010). In all crosses, alvear terminals were excited with 5 Hz optogenetic theta burst stimulation either without a concurrent orange light flash or with an orange light flash to silence PV interneurons (3000 ms duration) (Fig. 7). In mice without Arch-GFP containing PV cells, orange light had no effect on the IPSC amplitudes (Fig. 7A, n = 6). In contrast, orange light could completely suppress theta burst generated IPSCs in mice expressing Arch (Fig. 7B, n = 6). When we examined normalized IPSC amplitudes, orange light significantly suppressed IPSC amplitudes in mice that expressed Arch in PV interneurons (Fig. 7B, C, *P* < 0.0001 two-way ANOVA, Bonferroni post hoc test). The average magnitude of the orange light suppression in Arch-GFP PV interneurons was greater than 50% (orange light = 41.2 +/- 3.8% control amplitudes). Therefore, these interneurons contributed significantly to alvear pathway feedforward inhibition of CA1 pyramidal neurons in our mouse models.

**Figure 7.**
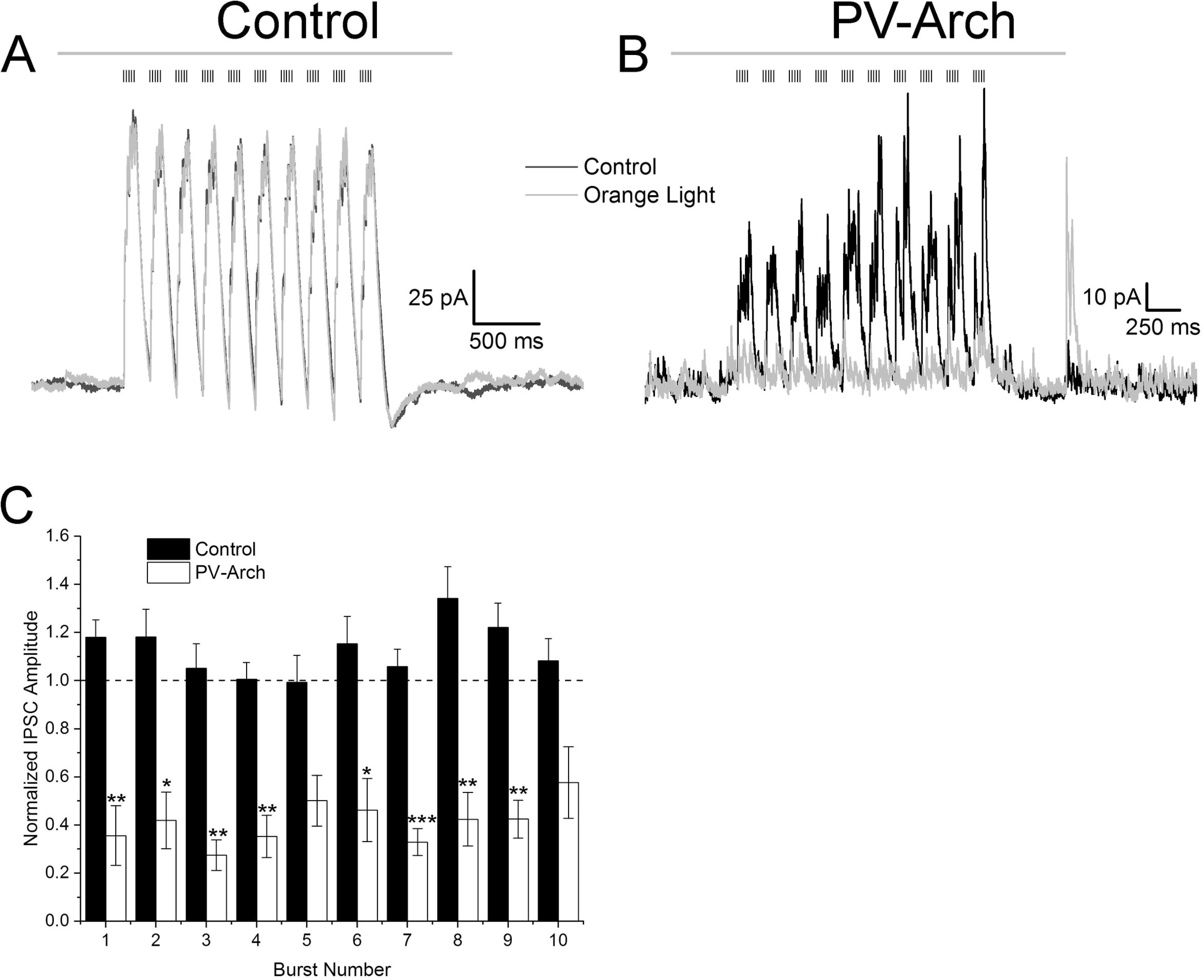
Suppression of PV interneuron activity inhibits alvear pathway driven disynaptic inhibitory currents in hippocampal CA1 pyramidal neurons. **A.** 5 Hz optogenetic theta burst (black vertical lines above traces) stimulation of alvear inputs produced outward IPSCs in a voltage clamped CA1 pyramidal neuron held at +15 mV. On alternating trials, an orange light pulse (grey line above traces, 3000 ms duration) was initiated 500 ms before and ended 620 ms after the theta burst stimuli. **B**. Same experimental design was used as A except Arch-GFP was expressed in PV interneurons. **C**. Bar plot illustrating the normalized maximum IPSC amplitude for each burst (Orange light/No Orange light). Black bars (control) were measured in pyramidal cells in mice lacking Arch-GFP whereas open bars were measured in pyramidal cells in PV;Arch-GFP mice (* *P* < 0.05, ** *P* < 0.01, *** *P* < 0.001).

## Discussion

Our results suggest that alvear inputs from the MEC were capable of producing facilitating low probability release glutamatergic synapses onto both PCs and interneurons in the SO of hippocampal CA1. The alvear inputs onto the basal dendrites of CA1 PCs produced subthreshold EPSPs even under robust theta burst stimulation paradigms. In contrast, theta burst stimulation of alvear inputs strongly activated horizontally oriented interneurons in the SO of CA1 producing suprathreshold responses. However, not all interneuron subtypes were equally affected by alvear inputs. PV perisomatic interneurons responded more robustly to alvear theta burst stimulation by producing higher frequencies of action potentials compared to VIP and GIN interneurons. Furthermore, optogenetic silencing of PV interneurons during alvear theta burst stimulation suppressed disynaptic rhythmic IPSCs in CA1 PCs by greater than 50%. Therefore, MEC alvear inputs appeared to primarily influence hippocampal CA1 via feedforward inhibition. This pathway may contribute to the generation of rhythmic perisomatic inhibition observed during theta rhythms in hippocampal CA1 PCs (Soltesz I and M Deschenes 1993).

In contrast to the alvear pathway, the EC TA pathway synapses on the distal apical dendrites of CA1 PCs in the SLM (Steward O 1976; Steward O and SA Scoville 1976). There is some disagreement on whether stimulation of the EC *in vivo* is capable of eliciting suprathreshold excitation of CA1 PCs (Andersen P *et al*. 1966; Segal M 1972; Yeckel MF and TW Berger 1990; Leung LS *et al*. 1995; Yeckel MF and TW Berger 1995). However, *ex vivo* stimulation of the TA or direct activation of the SLM has consistently been found to result in subthreshold EPSPs in CA1 PCs (Colbert CM and WB Levy 1992; Empson RM and U Heinemann 1995, 1995; Ang CW *et al*. 2005). Furthermore, in the absence of synaptic inhibition, stimulation of the SLM produced facilitating EPSPs suggestive of a low probability release synapse (McQuiston AR 2007, 2008). Thus, similar to findings in the TA, stimulation of MEC alvear terminals resulted in facilitating EPSCs indicative of a low probability release synapse.

Additionally, MEC alvear inputs onto CA1 PCs appear to be monosynaptic because TTX did not completely eliminate EPSCs in CA1 PCs, and the potassium channel blocker 4-AP partially rescued the response to alvear input. Furthermore, MEC alvear excitatory inputs onto CA1 PCs were mediated by AMPA receptor activation. In contrast, TTX completely inhibited MEC alvear input driven IPSCs in CA1 PCs. Importantly, TTX inhibited IPSCs in CA1 PCs could not be rescued by 4-AP. These results suggest that inhibitory synaptic transmission driven by MEC alvear input stimulation was not monosynaptic. Furthermore, IPSCs in CA1 PCs driven by alvear input were blocked by both AMPA receptor antagonists and GABA_A_ receptor antagonists. These findings suggest that MEC alvear inputs drive disynaptic inhibition in hippocampal CA1 PCs. Therefore, although alvear inputs monosynaptically excite CA1 PCs, the predominant effect of MEC alvear inputs on CA1 PC activity may be through disynaptic inhibition.

Consistent with alvear disynaptic inhibition of CA1 PCs, activation of MEC alvear inputs produced EPSCs in CA1 SO interneurons. These EPSCs were facilitating, suggesting that MEC alvear inputs onto interneurons had a low probability of release. Furthermore, the MEC alvear inputs were mediated by AMPA receptors. Post hoc anatomical reconstruction of these CA1 SO interneurons showed that they had horizontal somatodendritic morphology, some of which had axons confined to the PC cell body layer. Given that these interneurons were confined to SO, alvear pathway activation must arise from activation of postsynaptic elements in SO and not due to the potential propagation of action potentials to alvear terminals located in the SLM.

Although the timing of action potentials in individual MEC L3 projection neurons do not correlate well with population EC theta field potentials, MEC L3 projection neurons do fire coherently with their own internal theta activity (Quilichini P *et al*. 2010; Domnisoru C *et al*. 2013). Indeed, theta burst stimulation of the SLM occasionally resulted in the generation of a distal apical dendritic spikes in CA1 PCs (Takahashi H and JC Magee 2009). However, robust optogenetic theta burst stimulation of the alvear pathway in the SO did not produce suprathreshold EPSPs measured from CA1 PC somas. These data may seem to suggest that the TA pathway input is stronger than the alvear pathway in producing responses in hippocampal CA1 PCs. However, stimulation in the SLM (Takahashi H and JC Magee 2009) would activate more than just TA synaptic terminals. SLM stimulation potentially activates alvear inputs that ultimately project to the SLM (Deller T *et al*. 1996) as well as glutamatergic inputs that arise from the thalamic nucleus reuniens (Wouterlood FG et al. 1990; Dolleman-Van Der Weel MJ and MP Witter 1996). Therefore, a direct comparison between the strengths of TA and alvear inputs was not possible with previous studies. Nevertheless, our studies do suggest that stimulation of SO alvear inputs were not strong enough to drive CA1 PCs to fire action potentials.

In contrast to CA1 PCs, theta burst stimulation of the MEC alvear pathway was capable of eliciting suprathreshold excitation of interneurons located in the SO and SP of hippocampal CA1. However, not all subtypes of interneurons were equally excited by theta burst stimulation of the alvear pathway. Interneurons expressing PV produced a higher frequency of action potentials relative to interneurons expressing VIP or GIN interneurons. Furthermore, a larger proportion of PV and VIP interneurons produced suprathreshold or subthreshold responses compared to GIN interneurons. Importantly, different subtypes of PV and VIP interneurons could produce bursts of action potential by theta burst stimulation of the alvear pathway. These responsive interneuron subtypes included PV and VIP perisomatic interneurons as well as PV bistratified and presumptive VIP interneuron-selective interneurons (Booker SA and I Vida 2018). Importantly, PV interneurons have been demonstrated to produce little to no response to SLM stimulation (Milstein AD *et al*. 2015). Therefore, alvear pathway terminals in the SO appear to be capable of activating different subsets of interneurons relative to TA inputs.

PV interneurons play a crucial role in coordinating biologically relevant rhythms in populations of hippocampal principal cells (Buzsaki G and XJ Wang 2012). These rhythms include gamma and theta rhythms. Our observations that alvear theta burst stimulation produced rhythmic IPSCs in CA1 PCs is consistent with previous observations. Moreover, we have also demonstrated that theta burst stimulation of the alvear pathway can generate burst firing of PV interneurons. Importantly, optogenetic silencing of PV interneurons during alvear theta burst stimulation resulted in a greater than 50% reduction in burst IPSC amplitudes observed in CA1 PCs. Given that the largest source of theta rhythm generation arises from the EC (Buzsaki G et al. 1983; Bragin A *et al*. 1995; Kamondi A *et al*. 1998), our data suggest that alvear excitation of PV interneurons in hippocampal CA1 may contribute to theta rhythm generation by perisomatic interneurons (Buzsaki G 2002; Buzsaki G and XJ Wang 2012).

In conclusion, our study demonstrated that the alvear pathway from the MEC innervated both CA1 PCs and interneurons. These glutamatergic excitatory inputs were weak and incapable of activating CA1 PCs but were capable of eliciting action potentials in interneurons. However, not all interneurons were equally affected. PV-expressing interneurons were more excitable compared to VIP-expressing and GIN interneurons (a subset of O-LM interneurons). Greater than half of the feedforward inhibition in CA1 PCs produced by theta stimulation of the alvear pathway was due to the activation of PV interneurons, many of which were perisomatic. Furthermore, the subset of interneurons activated by the alvear pathway in the deep layers of hippocampal CA1 appear to be different from those activated by the TA. Therefore, our data suggests that the MEC alvear pathway primarily affects hippocampal CA1 function by driving feedforward inhibition. Furthermore, we propose that this pathway may contribute to the generation of theta rhythms.

## Acknowledgements

The authors would like to thank Dr. Susumu Tonegawa for generously donating the pOXR-1-Cre mice, Dr. Leonardo Belluscio for donating the tetO-oChIEF-mCitrine mice, and Dr. John Dempster for providing WCP electrophysiological software. This work was supported by funding from the National Institutes of Health (R01MH107507 and R21AG055073 to ARM).

